# Structural characterization of a MAPR-related archaeal cytochrome b_5M_ protein

**DOI:** 10.1101/2021.11.30.470528

**Authors:** Sarah Teakel, Michealla Marama, David Aragão, Sofiya Tsimbalyuk, Jade K. Forwood, Michael A. Cahill

## Abstract

We recently reported that the membrane associated progesterone receptor (MAPR) protein family (mammalian members: PGRMC1, PGRMC2, NEUFC and NENF) originated from a new class of prokaryotic cytochrome b_5_ (cytb_5_) domain proteins, called cytb_5<u>M</u>_ (<u>M</u>APR-like). Relative to classical cytb_5_ proteins, MAPR and ctyb_5M_ proteins shared unique sequence elements and a distinct heme binding orientation at an approximately 90⁰ rotation relative to classical cytb_5_, as demonstrated in the archetypal crystal structure of a cytb_5M_ protein (PDB accession number 6NZX). Here, we present the second crystal structure of an archaeal cytb_5M_ domain (*Methanococcoides burtonii* WP_011499504.1, PDB:6VZ6). It exhibits similar heme-binding to the 6NZX cytb_5M_, supporting the deduction that MAPR-like heme orientation was inherited from the prokaryotic ancestor of the original eukaryotic MAPR gene.

## Introduction

We recently investigated a possible eukaryogenic role of MAPR proteins in mitochondrial origins [1], prompted by the discovery that some mitochondrial genes had co-evolved with PGRMC1 [2], and that introducing point mutations at known phosphorylation sites on PGRMC1 had an effect on mitochondrial shape and mitochondrial protein abundance [3]. Several PGRMC1 functions also appear to be ancient in eukaryotes [1], including regulation of heme synthesis [4], cytP450 interactions [5] and sterol metabolism [2]. That investigation led to the discovery of the newly identified yet ancient cytb_5M_ sub-class of cytb_5_-domain proteins, which were more similar to MAPR than to classical eukaryotic cytb_5_ proteins and which therefore gave rise to MAPR proteins [1].

PGRMC1 is the archetypal and best characterized member of the heme binding eukaryotic MAPR family [1, 6] in which the heme-interacting tyrosine residue is Y113, and residues Y107, K163 and Y164 hydrogen bond (H-bond) with heme [7]. A group of cytb_5M_-like proteins from candidate phyla radiation (CPR) bacteria appear to exhibit similar tyrosinate heme chelation to MAPR proteins, because polar heme-interacting residues are strongly conserved with PGRMC1. We called these <u>tyrosine-(Y)</u>-containing cytb_5M_-like proteins cytb_5MY_. It has so far not been possible to determine whether MAPR proteins evolved from a CPR cytb_5MY_, or whether cytb_5MY_ arose after horizontal gene transfer of a MAPR gene into CPR bacteria [1].

While the prokaryotic cytb_5M_ and cytb_5MY_ proteins share a conserved surface patch and heme-binding pocket similarities with MAPR proteins, they lack a MAPR-specific inter-helical insertion region (MIHIR) sequence that is found in MAPR proteins. The PGRMC1 MIHIR extends from L130 to E157 in the protein sequence, between residues that H-bond with heme, but which loop away to the opposite protein surface from the heme-binding site in the folded protein structure [1, 7]. A conserved motif in the MIHIR region of PGRMC1 resembles a coiled-coil motif found in several myosin proteins, suggesting that PGRMC1 may interact with some of the same proteins as myosins, i.e. components of the actin cytoskeleton [6]. PGRMC1 was co-immunoprecipitated with actin cytoskeleton associated proteins including RACK1 and α-Actinin-1 [8, 9]. Therefore, the MIHIR, a eukaryotic invention, may be involved with actin cytoskeletal interactions. Point mutations at known phosphorylation sites on PGRMC1, including tyrosine residue 180 affected cell motility and altered the protein abundance of actin cytoskeleton associated proteins [3, 10]. It is probable that interactions between PGRMC1 and components of the actin cytoskeleton, predicted to occur within the previously described MIHIR sequence, could be regulated by tyrosine phosphorylation [6].

In this present study we solved the crystal structure of a second cytb_5M_ protein, which demonstrated substantial overall three-dimensional similarity to our previous cytb_5M_ structure [1]. These results will contribute to our future characterization of MAPR origins, and the understanding of how eukaryotes adapted ancient functions and invented novel ones to produce the first MAPR protein, whose strongly conserved inheritance into phylogenetically diverse eukaryotes suggests it performed essential roles in a very early eukaryote [1, 6].

## Methods

The archaeal *Methanococcoides burtonii* WP_011499504.1 protein selected for this study was previously identified as a cytb_5M_ protein [1]. To produce pure protein to determine the structure by protein crystallization protein constructs were expressed and purified in *Escherichia coli PLysS* cells as previously described [1]. Briefly, the codon-optimized WP_011499504.1 ORF was subcloned into pGEX-4T-1-H expression vector (Genscript) to create pGEX4T1_ WP011499504. Competent *E.coli BL21(DE3) PLysS* cells (Novagen #69451) (50 μL) were transformed with 1 μL of plasmid DNA using heat shock method and cell recovery. 80 μL of cells were spread onto Luria Base plates containing ampicillin (100 ug/mL) and incubated overnight at 37°C. Luria Broth containing ampicillin (100 ug/mL) was inoculated with the bacterial colonies and incubated overnight at room temperature at 220 rpm. MiniPrep (Qiagen) was performed using the transformed *E.coli* cells as per manufacturer’s protocol. Large scale cultures were produced using the transformed *E.coli* cells by adding 500 μL of cells to 500 mL of autoinduction expression base media (1 % Tryptone, 0.5 % yeast extract, 1 mM MgSO4, 5 % 20x NPS, 2 % 50x 5052) containing ampicillin (100μg/ml). Cells were incubated at 30°C at 80 rpm for approximately 28 hours. To perform GST affinity chromatography using fast protein liquid chromatography (FPLC), the soluble cell extracts were injected using a superloop at 2 mL/minute into a GST column equilibrated with GST cell lysis buffer. The column was washed with 10 column volumes of GST buffer (50 mM Tris(hydroxymethyl)aminomethane, 125 mM NaCl, pH 7.4). Competitive binding using GST buffer containing 10 mM Glutathione eluted purified GST-tagged protein from the column. To cleave the N-terminal GST tag from the protein, 100 μL of Tobacco etch virus (TEV) protease was added to the protein eluate and incubated at 4°C overnight with TEV. Size exclusion chromatography was performed using AKTA FPLC with S200 20/60 filtration column equilibrated with 50 mM Tris. The purity of the samples and the complete cleavage of the cytb_5M_ domain from the affinity tag were assessed using SDS PAGE analysis.

Diffractable protein crystals were produced in an optimised condition from the PEG ION 2 screen (Hampton Research) using purified protein at 8 mg/mL in a condition containing 0.1 M succinic acid, 12 % PEG6000 incubated at 23°C. Crystals produced from a crystal optimisation were flash frozen using liquid nitrogen in a 20 % glycerol stock and sent for x-ray diffraction to the Australian Synchrotron Facility, Melbourne. Crystal data was collected using the MX2 (Eiger 16M detector) crystal beamline [11, 12] and Blu-Ice software [13] at the Australian Synchrotron Facility. The data was integrated in iMosflm [14] scaled and reduced in AIMLESS [15]. The structure was determined by molecular replacement using PDB ID 1J03 in Phaser [16], REFMAC [17], PHENIX [18] and COOT [19].

## Results

### *Methanococcoides burtonii* WP_011499504.1 cytb_5M_ protein purification

The archaeal *M. burtonii* cytb_5M_ domain was recombinantly expressed and purified (Figure 1) prior to successful crystallization and structural comparisons with the crystal structure of the cytb_5_ domains of PGRMC1 (PDB: 4X8Y) [7] and our previously described *Hadesarchaea* cytb_5M_ protein structure (PDB: 6NZX) [1]. Both cytb_5M_ proteins are from the archaeal Euryarchaeota taxon. The *M. burtonii* cytb_5M_ domain existed predominantly as an ~8 kDa monomer in solution on S200 gel filtration profile. A peak at ~50 kDa is also present, representing a dimer of the GST affinity tag (Figure 1).

**Figure 1.**
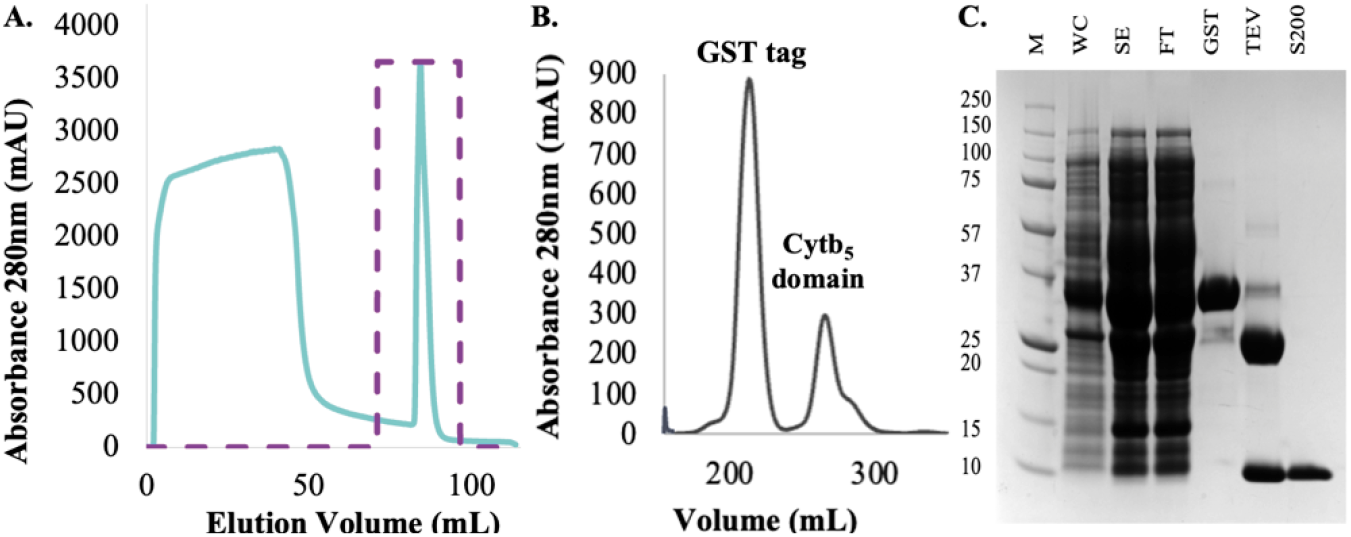
*Methanococcoides burtonii* WP_011499504.1 cytb_5M_ domain expression and purification. **A.** GST affinity purification of the *M. burtonii* cytb_5M_ domain UV trace (blue) and fractions containing purified protein (purple) **B.** S200 size exclusion chromatography of the cytb_5M_ domain. **C.** SDS PAGE analysis showing a purified protein band at ~8 kDa. **Legend: M** Marker **WC** Whole cell extract **SE** Soluble extract **FT** Flow through **GST** GST-tagged purified protein (~33kDa) **TEV** Cleavage of GST tag (~25kDa) with TEV protease from purified protein (~8kDa) **S200** Purified protein (~8kDa)

The crystal structure of the *Methanococcoides burtonii* WP_011499504.1 cytb_5M_ domain Diffractable protein crystals were produced in an optimized condition from the PEG ION 2 screen (Hampton Research) using purified *M. burtonii* cytb_5M_ domain protein at 8 mg/mL in a condition containing 0.1 M succinic acid, 12% PEG6000 incubated at 23 °C (Figure 2a). The structure of the *M. burtonii* cytb_5M_ domain was solved to a resolution of 2.1 Å (Table 1) and contained a bound cofactor, identified as heme (Figure 2b). This is consistent with the red coloration of the protein in solution and of the protein crystals (Figure 2a) and was also evidenced through absorbance at ~412 nm using spectroscopy (not shown). Although the heme is not covalently bonded in the structure of the *M. burtonii* cytb_5M_ domain, it is buried within the protein (Figure 2c).

**Table 1.** Crystallography data collection and refinement statistics

|  |  |
| --- | --- |
| Resolution range | 67.5-2.1 |
| Space group | P 4 2 2 |
| Unit cell | 67.495 67.495 48.132 90 90 90 |
| Reflections | 6908 |
| Multiplicity | 7.1 (4.8) |
| Completeness (%) | 98.0 (95.0) |
| Mean I/sigma(I) | 14.3 (5.1) |
| R-merge | 0.089 (0.297) |
| R-meas | 0.096 (0.334) |
| R-pim | 0.033 (0.149) |
| CC1/2 | 0.997 (0.946) |
| Reflections used in refinement | 6717 (631) |
| R-work | 0.1622 |
| R-free | 0.1953 |
| Number of non-hydrogen atoms | 726 |
| macromolecules | 617 |
| ligands | 43 |
| solvent | 66 |
| Protein residues | 79 |
| RMS (bonds) | 0.0102 |
| RMS (angles) | 1.04 |
| Ramachandran favoured (%) | 98.67 |
| Ramachandran allowed (%) | 1.33 |
| Ramachandran outliers (%) | 0.00 |
| Rotamer outliers (%) | 0.00 |
| Clashscore | 4.84 |
| Average B-factor | 24.8 |
| macromolecules | 24.2 |
| ligands | 22.1 |
| solvent | 32.7 |

Statistics for the highest-resolution shell are shown in parentheses.

**Figure 2.**
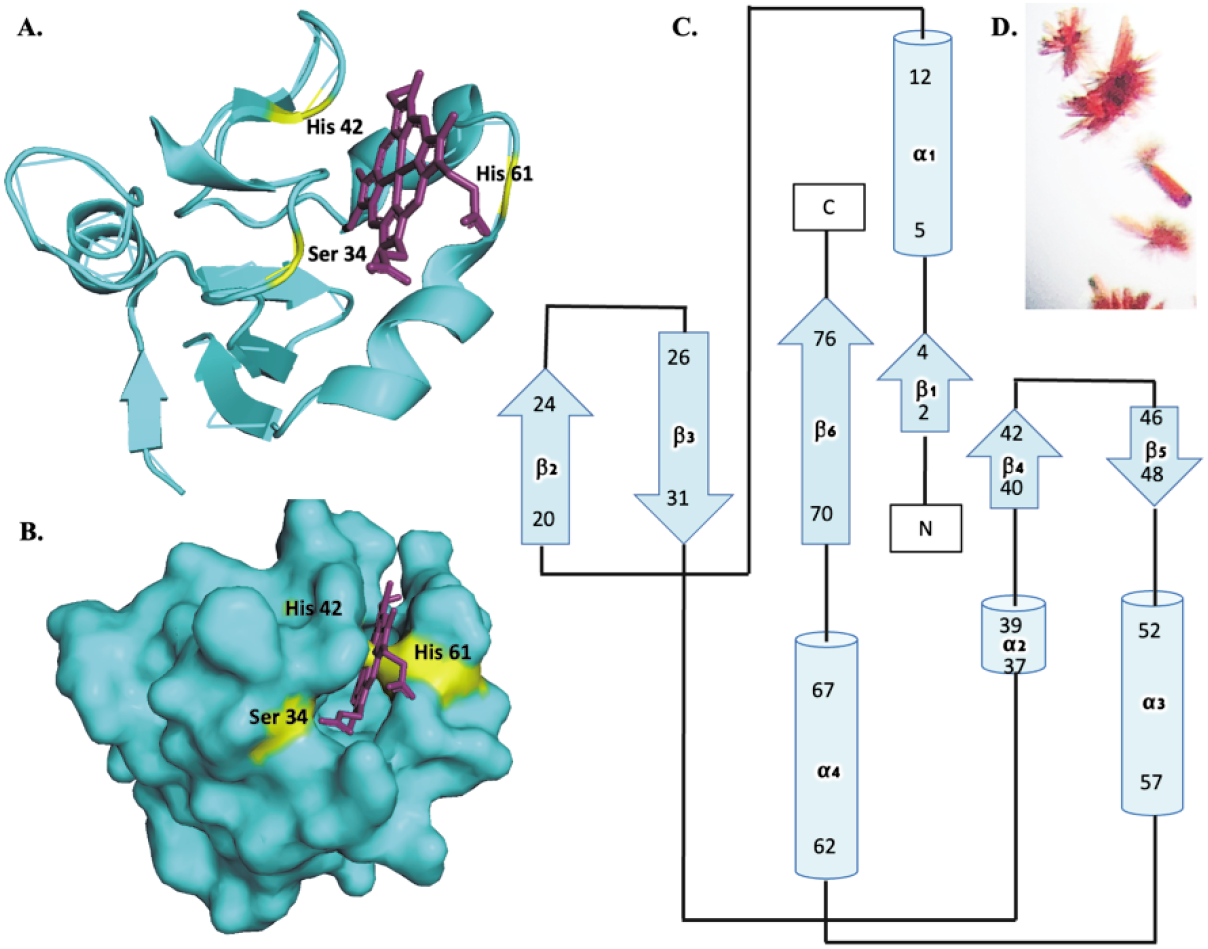
Schematic representation of the structure of the *Methanococcoides burtonii* WP_011499504.1 cytb_5M_ domain. **A.** The structure of the cytb_5M_ domain (blue) showing a heme bound ligand (purple). **B.** Surface representation (blue) and the heme ligand (purple), which is buried within the cytb_5M_ domain. Key interacting residues (S3, H42 and H61) are shown (yellow). **C.** Crystals were obtained in 0.1M succinic acid, 12% PEG6000 at 23°C. The red coloration is due to the presence of the heme ligand. **D.** Topology map obtained from PDBsum showing alpha helices (cylindrical) and beta sheets.

Diffraction data was indexed and integrated in the space group P 4 2 2, with unit cell dimensions of *a* =67.495=, *b* = 67.495, and *c* = 48.132, and angles of *a* = 90, *b* = 90, and *c* = 90. Following rebuilding and refinement in COOT and REFMAC respectively, the final model had an *Rwork* and *Rfree* of 0.1622 and 0.1953 respectively, no Ramachandran outliers, and good stereochemistry (Table 1). The structure was analyzed using Proteins, Interfaces, Structures and Assemblies (PISA) [20]. The protein and ligand complex had an interface area of 524.4 Å (Figure 2c). The coordinates and associated structural data were deposited and validated to the Protein Data Bank (PDB) and issued the code 6VZ6.

As predicted from the protein sequences, there is high structural similarity between the *M. burtonii* and the *H. archaeon ynp_n21* cytb_5M_ domains (Figure 3a). The main polar interactions with heme include the side chain S34 (Figure 3b). The H-bond interactions are shown in Figure 3e. The structure of *H. archaeon ynp_n21* cytb_5M_ domain showed heme chelation through residues H42 and H61 (Figure 3c). The cognate residues were involved in H-bonding in *M. burtonii* cytb_5M_ domain (Figure 3b). Differences in the residues that interact with heme between cytb_5M_ and MAPR proteins indicate the appearance of novel heme-binding properties in the evolutionary transition from prokaryotic cytb_5M_ to eukaryotic MAPR proteins, as exemplified by PGRMC1 (4X8Y [7]).

**Figure 3.**
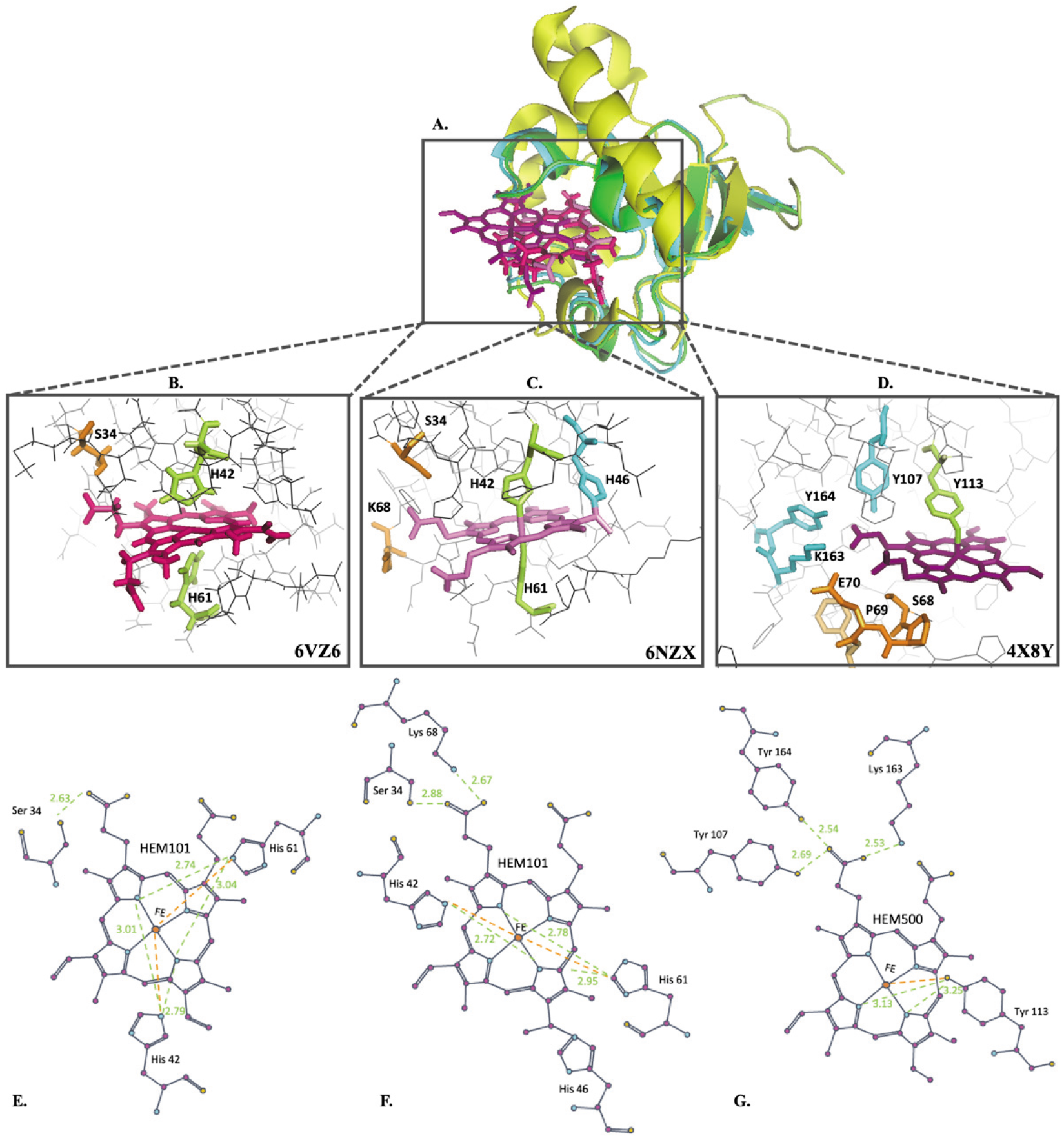
Heme axial binding within the cytb_5M_ domain. **a.** Overlay of archaeal cytb_5M_ domains and PGRMC1 cytb_5_/MAPR domain structures from PDB. The PGRMC1 cytb_5_ domain (yellow) has a longer helix than the *H. archaeon ynp_n2* and *M. burtonii* cytb_5M_ domains (blue and green respectively), which is shifted away from the heme molecule. **b.** Heme binding within the *M. burtonii* WP_011499504.1 cytb_5M_ domain involves H-bonding at S34 (orange) and heme Fe axial ligand support at H42 and H61 (green). **c.** Heme binding within the *H. archaeon ynp_n21* cytb_5M_ domain involves binding at H46 (blue), H-bonding at S34 and K68 residues (orange) and heme axial ligand support at H42 and H61 (green). **d**. Heme binding within the PGRMC1 cytb_5_/MAPR domain is coordinated through Y113 residue (green) and H-bonding at Y107, K163 and Y164 (blue). Note the absence of Fe-interacting axial histidine residues, indicating that this type of heme interaction differs from that seen for the archaeal cytb_5M_ proteins. **Figures e-g.** are adapted from figures produced using LIGPLOT [25]. Note the residues in orange (S68, P69 and E70) are interactions from the bacterial expression vector with the heme, not PGRMC1 residues, as previously discussed [1]. **e.** Heme binding in the *M. burtonii* WP_011499504.1 cytb_5M_ domain. **f.** Heme binding in the *H. archaeon ynp_n21* cytb_5M_ domain. **g.** Heme binding within the cytb_5_/MAPR domain of PGRMC1.

A heuristic PDB search was performed using DALI protein structure comparison by alignment of distance matrices [21] to identify the protein structures, from those available on the PDB, with the highest structural homology to the *M. burtonii* cytb_5M_ domain (Table 2). As expected, the *M. burtonii* cytb_5M_ domain showed highest homology to the *H. archaeon ynp_n21* cytb_5M_ domain with a root mean square deviation (RMSD) of 0.7 and %ID of 54%. The protein identified with the second highest confidence score was the PGRMC1 cytb_5_/MAPR domain, although comparatively the similarity was much lower, with an RMSD of 2.2 and %ID of 23%.

**Table 2.** Proteins with the highest structural homology to archaeal *Methanococcoides burtonii* cytb_5M_ domain (6VZ6)

| PDB ID | z | RMSD | LALI | Nres | %ID | Protein name | Gene | Organism |
| --- | --- | --- | --- | --- | --- | --- | --- | --- |
| 6NZX | 16.4 | 0.7 | 76 | 76 | 54 | Cytochrome b5 | APU95_03975 | *H. archaeon YNP_N21* |
| 4X8Y | 8.3 | 2.2 | 73 | 112 | 23 | Progesterone receptor membrane component 1 | PGRMC1 | *Homo sapiens* |
| 1SOX | 8.2 | 2.0 | 69 | 463 | 25 | Sulfite oxidase | SUOX | *Gallus gallus* |
| 1X3X | 8.0 | 2.4 | 71 | 82 | 25 | Cytochrome b5 | N/A | *Ascaris suum* |
| 1KBI | 5.9 | 2.4 | 67 | 504 | 30 | Cytochrome b2 | CYB2 | *Saccharomyces cerevisiae* |
| 2I96 | 5.8 | 2.8 | 69 | 108 | 25 | Cytochrome b5 | CyB5A | *Homo sapiens* |
| 2KEO | 2.6 | 3.6 | 46 | 92 | 28 | Probable E3 ubiquitin-protein ligase HERC2 | HERC2 | *Homo sapiens* |

Z- z-score confidence in similarity significance, RMSD- Root mean square deviation across the aligned sequences, LALI- Total number of aligned residues, Nres- Total number of residues, %ID- Percent identity.

All other identified sequences belong to the classical type cytb_5_ group, underlining the novelty of the cytb_5MY_ structural fold and its similarity to the only solved MAPR crystal structure, that of PGRMC1.

## Discussion

MAPR proteins originated from a prokaryotic cytb_5M_ domain protein, or possibly from the sub-class cytb_5MY_, rather than from the conventionally-recognized cytb_5_ proteins that gave rise to classical mammalian cytb_5_ proteins [1]. Here, we present the second structure of a cytb_5M_ protein, from *M burtonii* (WP_011499504.1). Unlike PGRMC1, the heme iron atom of both the archaeal *H. archaeon ynp_n21* and *M. burtonii* WP_011499504.1 cytb_5M_ domains are coordinated by two conserved histidine residues that serve as axial ligands (H42 and H61). The dual histidine mode of heme chelation (Figure 3) is conserved in cytb_5M_ domain proteins such as the previous 6NZX cytb_5M_ structure, as well as classical cytb_5_ domain-containing proteins [1]. The heme is buried within the cleft, further supported through conserved H-bonding at S34. This differs from the tyrosinatate Y113-mediated heme Fe chelation observed in PGRMC1 [7]. The situation in other MAPR proteins remains unknown but is presumed to resemble PGRMC1. Differences in heme binding are attributable to evolutionary modification of heme-interacting residues in PGRMC1 relative to the cytb_5M_ proteins (Figure 3, Figure 4).

**Figure 4.**
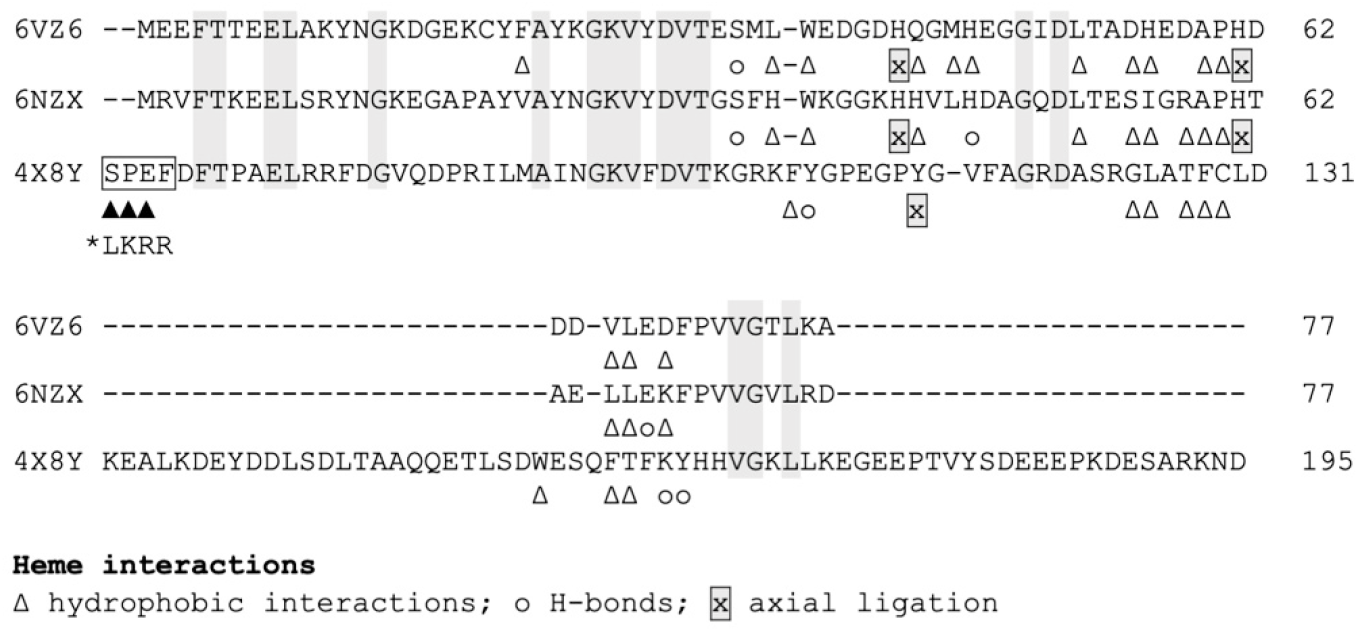
Sequence alignment of *M. burtonii* (6VZ6) and *H. archaeon ynp_n21* (6NZX) cytb_5M_ domains and PGRMC1 (4X8Y) cytb_5_ domain from a structural comparison using the *M. burtonii* cytb_5M_ domain protein sequence (6VZ6) as the reference. Residues conserved in all three proteins are highlighted in grey. Filled triangles (▲) at the N-terminus of 4×8Y represent heme interactions with non-PGRMC1 residues in 4X8Y that are contributed by the bacterial protein expression cloning vector (boxed). The asterisked PGRMC1 residues 67-71 show the wild-type PGRMC1 sequence. Numbering refers to PGRMC1 (O00264) residues. The gapped alignment of the top (6VZ6 1-62) is based on the alignment of 1620 cytb_5_ domain proteins of clades-1 and -2 from Tamarit et al. ([1], Fig.3A). The lower alignment (6VZ6 63-77) is generated here based upon heme interactions.

Neither of the archaeal cytb_5M_ proteins exhibited a heme-dependent dimerization, as reported for PGRMC1 [7]. We note heme-stacking interactions observed in the PGRMC1 crystal structure could also potentially facilitate even larger multimer formation. This merits future investigation. PGRMC1 dimers or higher order structures have been observed not only in the bacterially expressed crystallized protein [7], but also in mammalian cells [22], where an anti-FLAG tag antibody can immunoprecipitate FLAG-tagged as well as endogenous PGRMC1. Endogenous PGRMC1 precipitation was dependent upon heme availability. Also, the molecule glycyrrhizin (the active compound in liquorice), and derivatives, binds to PGRMC1 residues involved in the dimeric interface, disrupting the protein complex and favoring monomer formation [22].

Although the structure of a cytb_5MY_ protein could be informative, we have so far been unable to obtain one. Future determination of the structures of other MAPR proteins and cytb_5MY_ proteins will be required to determine whether tyrosinate heme-chelation is commonly accompanied by heme-mediated dimerization. The shared cytb_5M_/MAPR orientation of heme-binding is presumably related to an important MAPR-dependent eukaryotic function. In light of the current understanding of the structural differences between MAPR proteins, cytb_5M_, and classical cytb_5_ proteins, including MAPR tyrosinate coordinated heme binding, heme orientation, and the presence and function of the MIHIR binding region, future research should investigate the origins and function of MAPR proteins, with particular focus on the PGRMC1 membrane trafficking function [23, 24].

## Data accessibility

Data to support the findings of this study are openly available in the Protein Data Bank at https://www.rcsb.org/, reference number 6NZX.

## Acknowledgements

This research was supported by Charles Sturt University School of Biomedical Sciences (MAC) and PhD scholarship (ST) funding. This research was partly undertaken on the macromolecular crystallography beamlines at the Australian Synchrotron, part of ANSTO (DA).

## Author contributions

Conceptualization (MAC), designed and performed experiments (ST, MM, ST), x-ray diffraction and data collection (DA), data analysis (JKF, ST), manuscript preparation (ST, MAC), manuscript revision (JKF).

## Additional information

This research was undertaken in part using the MX2 beamline at the Australian Synchrotron, part of ANSTO, and made use of the Australian Cancer Research Foundation (ACRF) detector. The Full wwPDB X-ray Structure Validation Report for the resulting structure is provided as Supporting File 1.

## Notes

https://www.rcsb.org/structure/6VZ6

### Competing Interest Statement

The authors have declared no competing interest.

